# Relationship between Regional White Matter Hyperintensities and Alpha Oscillations in Older Adults

**DOI:** 10.1101/2020.09.04.283200

**Authors:** Deniz Kumral, Elena Cesnaite, Frauke Beyer, Simon M. Hofmann, Tilman Hensch, Christian Sander, Ulrich Hegerl, Stefan Haufe, Arno Villringer, A. Veronica Witte, Vadim Nikulin

**Affiliations:** Department of Neurology, Max Planck Institute for Human Cognitive and Brain Sciences, Leipzig, Germany; Institute of Psychology, Neuropsychology, University of Freiburg, Germany; Institute of Psychology, Department of Clinical Psychology and Psychotherapy, University of Freiburg, Germany; CRC Obesity Mechanisms, Subproject A1, University of Leipzig, Leipzig, Germany; Department of Psychiatry and Psychotherapy, University of Leipzig Medical Center, Leipzig, Germany; LIFE – Leipzig Research Center for Civilization Diseases, University of Leipzig, Leipzig, Germany; IUBH International University, Erfurt, Germany; Department of Psychiatry, Psychosomatics and Psychotherapy, Goethe University Frankfurt, Frankfurt, Germany; Berlin Center for Advanced Neuroimaging, Charité – Universitätsmedizin Berlin, Berlin, Germany; Bernstein Center for Computational Neuroscience Berlin, Berlin, Germany; Clinic of Cognitive Neurology, University Hospital Leipzig, Leipzig, Germany; Centre for Cognition and Decision Making, Institute of Cognitive Neuroscience, National Research University Higher School of Economics, Moscow, Russian Federation

**Author notes:** Corresponding Author: Deniz Kumral, and, Department of Neurology, Max Planck Institute for Human Cognitive and Brain Sciences, Leipzig, Germany, Stephan Str.1a, 04103, Leipzig, Germany. These authors contributed equally to the manuscript (Deniz Kumral, Elena Cesnaite).

**Keywords:** EEG, MRI, white matter hyperintensity, aging, alpha power, resting-state

## Abstract

Aging is associated with increased white matter hyperintensities (WMHs) and with the alterations of alpha oscillations (7–13 Hz). However, a crucial question remains, whether changes in alpha oscillations relate to aging per se or whether this relationship is mediated by age-related neuropathology like WMHs. Using a large cohort of cognitively healthy older adults (N=907, 60-80 years), we assessed relative alpha power, alpha peak frequency, and long-range temporal correlations (LRTC) from resting-state EEG. We further associated these parameters with voxel-wise WMHs from 3T MRI. We found that a higher prevalence of WMHs in the superior and posterior corona radiata as well as in the thalamic radiation was related to elevated alpha power, with the strongest association in the bilateral occipital cortex. In contrast, we observed no significant relation of the WMHs probability with alpha peak frequency and LRTC. Finally, higher age was associated with elevated alpha power via total WMH volume. Although an increase in alpha oscillations due to WMH can have a compensatory nature, we rather suggest that an elevated alpha power is a consequence of WMH affecting a spatial organization of alpha sources.

## 1. Introduction

White matter lesions, also known as white matter hyperintensities (WMHs), are highly prevalent in older adults and are of paramount clinical relevance since they are known to accompany cognitive decline and dementia (Birdsill et al., 2014; Debette and Markus, 2010; Habes et al., 2016). WMHs are considered to reflect mainly small vessel disease (Wardlaw et al., 2015), which typically affects periventricular regions and deep white matter sparing U-fibers (Habes et al., 2016). Little is known, however, whether and how WMHs impact functional measures of brain activity. Due to their location, WMHs may cause disconnection of neuronal populations (O’Sullivan et al., 2001). Theoretically, such damage of cortico-cortical and cortico-subcortical pathways is expected to alter the synchronized activity of neurons measured with M/EEG (Hindriks and van Putten, 2013).

One of the most prominent EEG rhythms are alpha oscillations (7-13 Hz), which have been shown to originate from thalamocortical and cortico-cortical interactions (Bazanova and Vernon, 2014; Lopes Da Silva et al., 1997). Importantly, measures of alpha oscillations have been related to many aspects of cognitive function (Klimesch, 1999) and also to endophenotypes of brain aging (Ishii et al., 2018; Knyazeva et al., 2018) either using alpha peak frequency or power. While individual alpha peak frequency has been consistently shown to decrease with age (Ishii et al., 2018; Knyazeva et al., 2018; Mierau et al., 2017), the findings on alpha power remain rather inconsistent. Previous EEG studies showed decreases of alpha power across the lifespan when using relatively large sample sizes (Babiloni et al., 2006a; Lodder and van Putten, 2011; Vysata et al., 2012): Yet these age-related reductions in alpha power were either not strongly present within the older age groups (>60 years of age; Lodder and van Putten, 2011) or not replicated (Sahoo et al., 2020; Scally et al., 2018).

Apart from these two measures of alpha oscillations, temporal dynamics of the signals can be quantified with auto-correlation showing to what extent a past of the signal relates to its future. A very slow attenuation of the auto-correlation, which can be described with a power law, is also referred to as long-range temporal correlations (LRTC). The presence of LRTC indicates scale-free properties of the signal fluctuation pattern that look similar at different time scales. LRTC in the amplitude envelope of the neuronal oscillations were shown to extend to tens or even hundreds of seconds (Linkenkaer-Hansen et al., 2001; Nikulin and Brismar, 2005). Importantly, the presence of LRTC is consistent with the idea that neuronal networks may operate at a critical state, characterized by a balance between inhibition and excitation (Linkenkaer-Hansen et al., 2001; Nikulin and Brismar, 2005; Palva et al., 2013; Shew and Plenz, 2013). LRTC exponent that represents the decay of the autocorrelation has been linked to functional connectivity measures (Zhigalov et al., 2017), brain maturation (Smit et al., 2011), and different aspects of cognition (Mahjoory et al., 2019; Samek et al., 2016; Smit et al., 2011). However, the link between LRTC and structural brain changes has not yet been examined.

As both static (i.e., power, individual alpha peak frequency) and dynamic (i.e., LRTC) measures of alpha oscillations might be affected by microstructural deteriorations, due to the disconnection among neural cells and damage to cortico-cortical and cortico-subcortical pathways (Madden et al., 2017), WMHs-associated alterations of EEG rhythms are plausible. However, there are only a few EEG studies that have directly investigated the relationship between alpha oscillations and WMHs or integrity (Babiloni et al., 2011, 2008a; Valdés-Hernández et al., 2010; van Straaten et al., 2012). Previously, local and global disturbances of brain anatomy like white matter microstructure (Hinault et al., 2020; Hindriks et al., 2015; Minami et al., 2020; Valdés-Hernández et al., 2010) have been found to be related to alpha rhythm affecting its peak frequency and power. For instance, a previous study with 222 subjects using Cuban Human Brain Mapping Project (Valdés-Hernández et al., 2010) provides evidence that alpha peak frequency can be associated with both decrease and increase (depending on the region) in the microstructure of thalamocortical or corticothalamic fibers assessed by Fractional Anisotropy (FA) using diffusion tensor imaging (DTI). Interestingly, so far only a few studies have investigated the relationship between alpha power and WMHs (Babiloni et al., 2009, 2008b, 2008a). For instance, it has been observed that higher alpha power was associated with higher scores of the prevalence of WMHs in individuals with mild cognitive impairment (Babiloni et al., 2008a). Similarly, a recent study (Quandt et al., 2020) reported that higher WHM lesion load was related to reduced EEG alpha connectivity measures in healthy older adults (N=35). However, to our knowledge, no link between voxel-wise whole-brain WMHs and different parameters of alpha oscillations has been investigated using a large sample of healthy older adults. Moreover, a crucial question still remains unresolved, for example, whether changes in alpha oscillations relate to normal aging *per se* or rather they represent the impact of age-related neuropathology, for instance, WMHs. In this study, using a large population-based sample, we investigated neurophysiological links between age, WMHs and alpha oscillations. More precisely, we investigated the association between age and parameters of alpha oscillations, and whether this relationship was mediated by WMHs. We further explored the association of WMHs with parameters of alpha oscillations in a topographically specific manner taking into account the location of the lesioned white matter tracts.

## 2. Methods

### 2.1. Participants

Participants were drawn from the population-based Leipzig Research Center for Civilization Diseases LIFE-Adult study (Loeffler et al., 2015). All participants provided written informed consent, and the study was approved by the ethics committee of the medical faculty at the University of Leipzig, Germany. The study was performed in agreement with the Declaration of Helsinki. A subset of participants underwent a 3-Tesla MRI head scan and resting-state (rs)EEG recordings on two separate assessment days. We selected participants above 60 years of age and without additional brain pathology or history of stroke, multiple sclerosis, epilepsy, Parkinson’s disease, intracranial hemorrhage, or brain tumors. We further excluded individuals whose rsEEG recordings were not temporally close to the MRI acquisition time and participants for whom alpha peak could not be identified. The details about the time differences between EEG and MRI measurement days can be found in Supplementary Figure 1 (*M* = 23.4 in absolute days). This resulted in a final sample of 907 participants (*M*= 69.49 ± 4.63 years of age, 380 female) for the rsEEG sensor space analysis. After excluding individuals with failed T1-weighted segmentation and head-modeling, the final sample for the rsEEG source analysis was 855 (*M*= 68.89 ± 4.66 years of age, 360 female). For a detailed overview of the selection process, see **Figure 1**.

**Figure 1.**
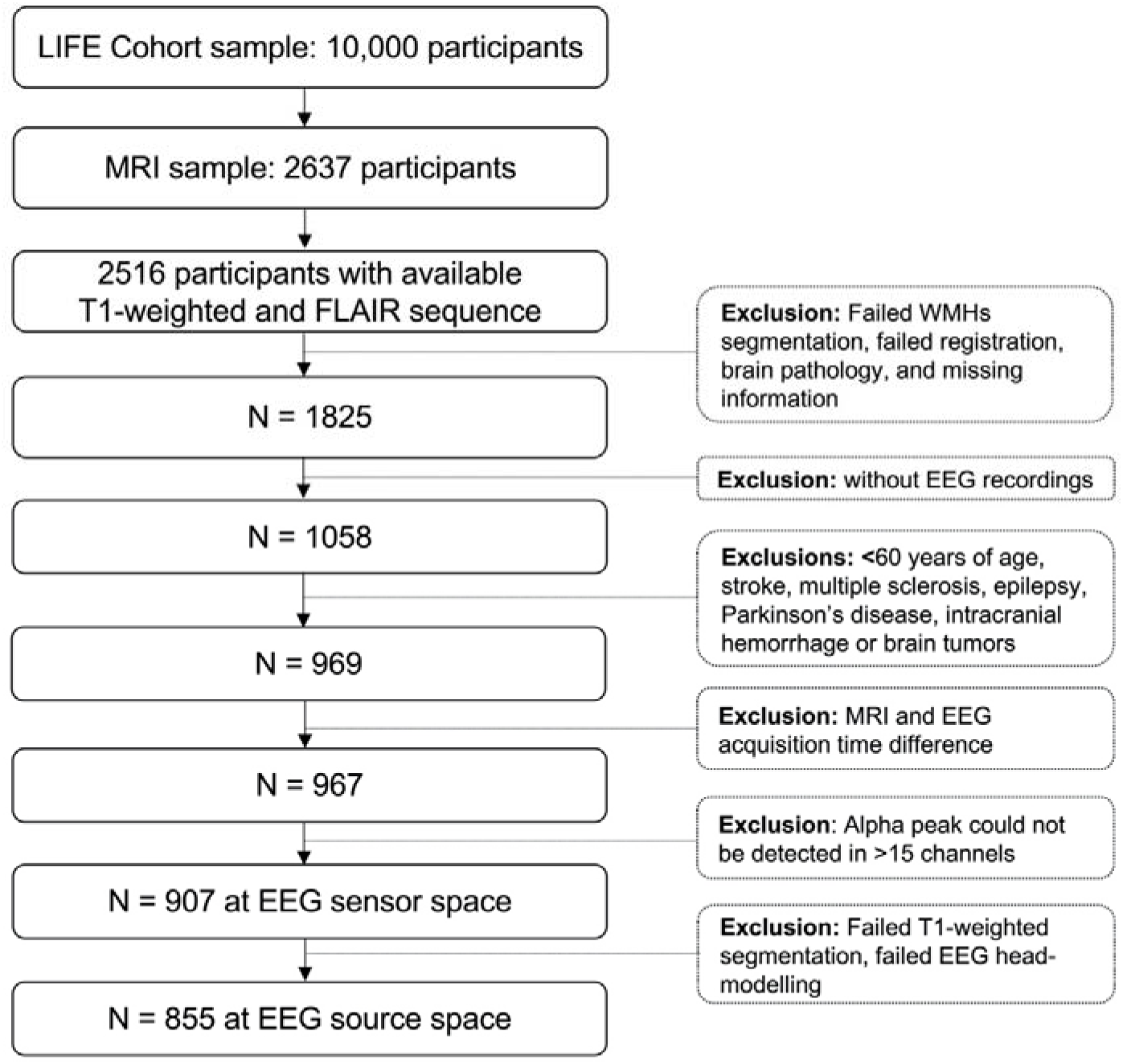
Flow chart visualizing the selection process of the MRI and EEG sample.

### 2.2. MRI Acquisition and Processing

All MRI scans were performed at 3 Tesla on a MAGNETOM Verio scanner (Siemens, Erlangen, Germany). The body coil was used for radiofrequency (RF) transmission and a 32-channel head coil was used for signal reception. T1-weighted MPRAGE and FLAIR images were acquired as part of a standardized protocol: MPRAGE (flip angle (FA) = 9°, relaxation time (TR) = 2300 ms, inversion time (TI) = 900 ms, echo time (TE) = 2.98 ms, 1-mm isotropic resolution, acquisition time (AT) = 5.10 min); FLAIR (TR = 5000 ms, TI = 1800 ms, TE = 395 ms, 1x0.49x0.49-mm resolution, AT = 7.02 min).

*Location of WMH.* The automated assessment of WMHs was computed in a previous study (Lampe et al., 2019). All images were checked by a study physician for incidental findings. A computer-based WMHs segmentation algorithm was then used to automatically determine WMH volume on T1-weighted MPRAGE and FLAIR images (Shiee et al., 2010) and inspected visually for segmentation errors. Binary WMH maps of all participants were nonlinearly co-registered to a standardized MNI template (1-mm isometric) with ANTS (Avants et al., 2011). In standard space, binary subject-wise WMH maps were grand-averaged to create a population WMH frequency map (Jenkinson et al., 2012) and to further compute the voxel-wise statistics. As previously implemented (Lampe et al., 2019), to segregate the periventricular (pv)WMH and deep (d)WMH, a default distance of 10 mm to the ventricular surface was used (DeCarli et al., 2005). Every voxel of WMH located within this border was classified as pvWMH; voxels outside the border were classified as dWMH.

*WMH Volume.* Regional WMH volume was calculated separately for the deep and periventricular white matter. Following Lampe et al. (2019), we added a constant value 1 to every participant’s regional dWMH volume because there were participants without lesions in the deep WM. We then calculated the ratio of dWMH and pvWMH (dWMH/pvWMH) as localized WMH volume. Total, deep and periventricular WMH volumes were further normalized to head size by total intracranial volume. Total and localized WMH (dWMH/pvWMH) volume were log-transformed for further statistical analyses.

### 2.3. EEG Acquisition and Preprocessing

RsEEG activity was recorded in an electrically and acoustically shielded room using an EEG cap with 34 passive Ag/AgCl electrodes (EasyCap, Brain Products GmbH, Germany). 31 scalp electrodes were placed according to the extended international 10–20 system. The signal was amplified using a QuickAmp amplifier, frequency range: DC-280 Hz (Brain Products GmbH, Germany). Two electrodes recorded vertical and horizontal eye movements while one bipolar electrode was used for electrocardiography. The rsEEG activity was referenced against common average and sampled at 1000 Hz. Impedances were kept below 10 kΩ. RsEEG data were preprocessed using EEGLAB toolbox (version 14.1.1b) and scripts were custom written in Matlab 9.3 (Mathworks, Natick, MA, USA). We filtered data between 1 and 45 Hz and applied a notch filter at 50 Hz. We then down-sampled the data to 500 Hz and ran a semi-automatic pipeline for artifact rejection: different noise threshold levels to mark bad time segments were used for the signal filtered in higher frequency (15–45 Hz) and lower frequency (1–15 Hz) ranges. The noise threshold for higher frequencies was set to 40 µV since noise at this range (i.e., induced by muscle activity) is typically lower in amplitude. The noise threshold for the lower frequency range was set to + 3SD over the mean amplitude of a filtered signal between 1 and 15 Hz. To control for the accuracy of automatically marked bad segments, we compared them to the noisy segments marked by another research group (Jawinski et al., 2017). Whenever these segments did not overlap by more than 10 s or they exceeded 60 s of total bad-segment duration, we inspected those datasets visually (∼10% of cases) to confirm whether they indeed were contaminated by noise. We further visually assessed power spectral densities (PSD) for data quality and used it to identify broken channels. Next, using independent component analysis (Infomax; Bell and Sejnowski, 1995), activity associated with the confounding sources — namely eye-movements, eye-blinks, muscle activity, and residual heart-related artifacts — was removed.

### 2.4. EEG Sensor Space Analysis

#### 2.4.1. Parameters of Alpha Oscillations

For rsEEG analysis, we used the first 10 min of a recording to avoid the potential effect of participants’ drowsiness. We individually adjusted the alpha band frequency range by locating a major peak between 7 and 13 Hz on Welch’s PSD with 4 s Hanning windows.

Thus, we determined individual alpha peak frequency in every channel and defined a bandwidth not exceeding 3 Hz around the peak. We then calculated relative alpha power for the individually adjusted alpha frequency range dividing it by the broadband power calculated in the 3–45 Hz frequency range. LRTC were calculated using detrended fluctuation analysis (DFA) on the amplitude envelope (calculated with Hilbert transform) of alpha band oscillations in time windows ranging from 3 to 50 seconds (while respecting the boundaries where the bad segments had been cut) based on the previously published procedure (Hardstone et al., 2012). Here, we briefly repeat the main steps: (1) a cumulative sum of the amplitude envelope is calculated, (2) the signal is then divided into pre-defined window sizes (τ) (3) the linear trend is removed in a given window. Fluctuation function F(τ) for all time windows of a given size τ is calculated as the root-mean-square of the detrended signal. In the case of a power-law relationship, we have F(τ) ∝ τ, where *v* is a scaling exponent (measuring LRTC) which can be obtained as a slope of a linear fit in log-log plot between F(τ) and τ. An exponent of 0.5 reflects uncorrelated signals (i.e., resembling white noise), *v*<0.5 indicates anticorrelations, while an exponent between 0.5<*v*<1 shows persistent autocorrelation (LRTC) where large fluctuations are likely to be followed by large fluctuation (Hardstone et al., 2012). This range of 0.5<*v*<1 is a typical range for many EEG and MEG studies.

The illustration of parameters of alpha oscillations are shown in **Figure 2**.

**Figure 2.**
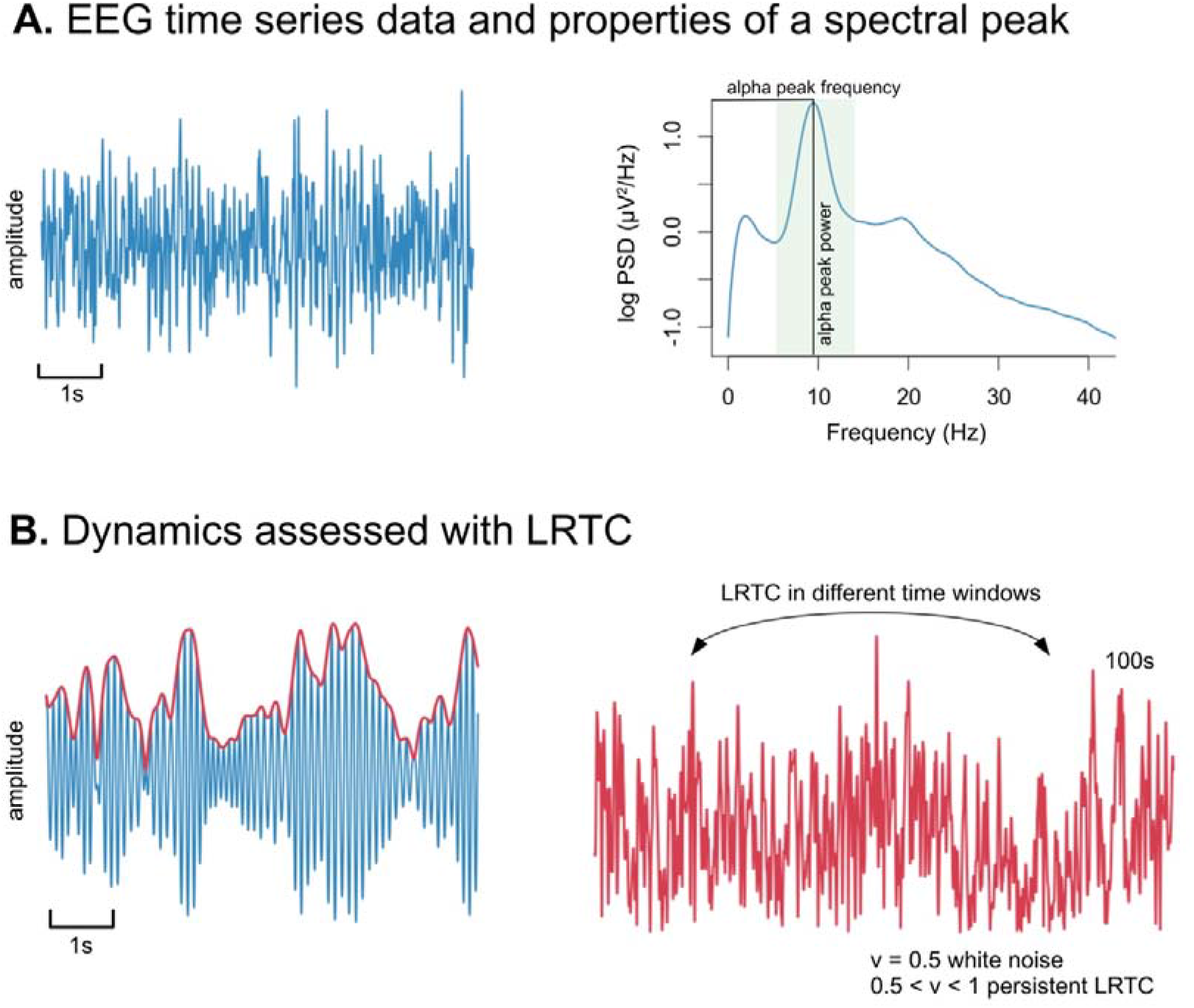
Illustration of parameters of alpha oscillations. **A)** Raw resting-state EEG time series data (blue) consists of various frequency bands that can be defined by their power and peak frequency. **B)** The temporal dynamics of a signal filtered in the alpha frequency range (8–12 Hz) is assessed by the properties of its amplitude envelope (red) using long-range temporal correlations (LRTC). The scaling exponent (ν) quantifies the presence of LRTC.

To reduce data dimensionality of rsEEG sensor space data used for the whole-brain voxel-wise inference analyses, we further grouped EEG channels into six coarser brain regions (frontal, central, temporal, parietal, and occipital) (**Figure 3A)**.

**Figure 3.**
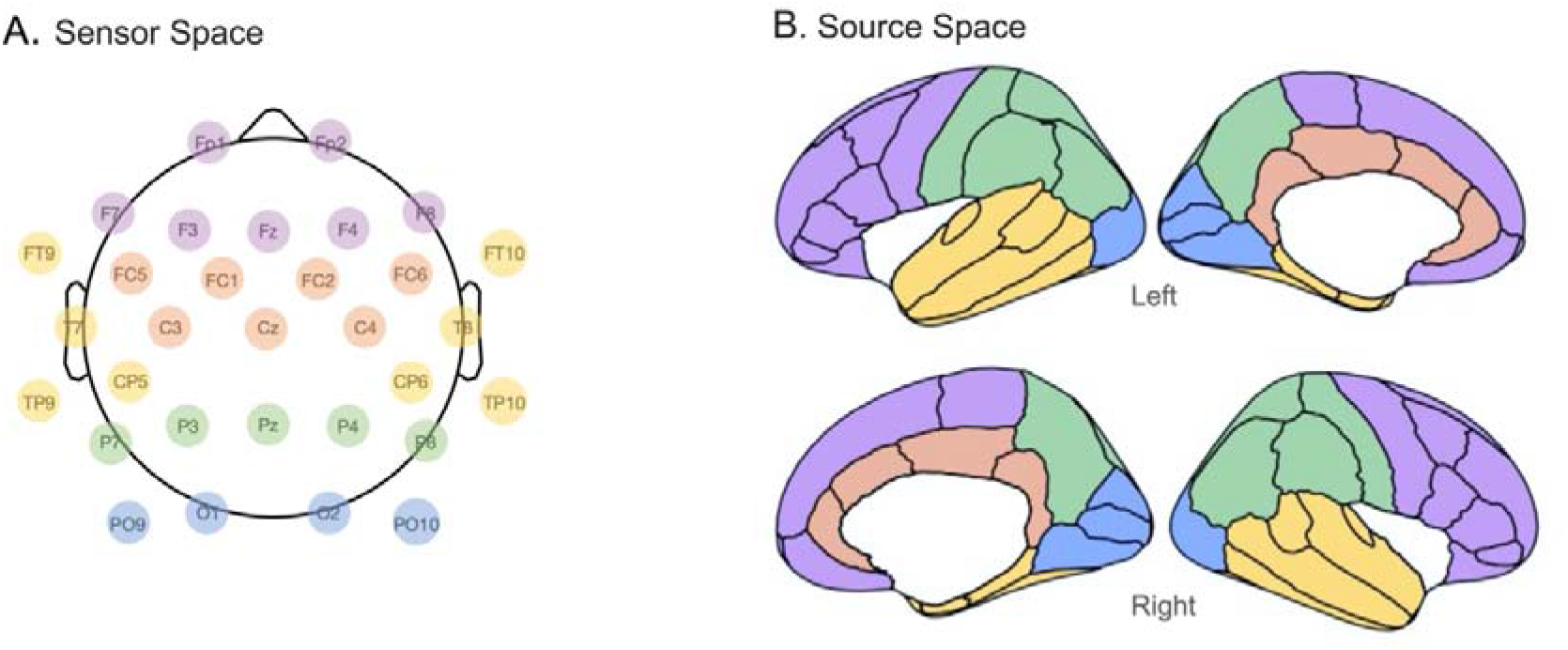
– Illustration of the regions of interest (ROIs) identified for EEG. Schematic topography for resting-state EEG in **A)** sensor space and **B)** source space. ROIs which form the frontal region are in purple, central region, and cingulate region (source) in orange, temporal region in yellow, parietal region in green, and occipital region in blue.

### 2.5. EEG Source Space Analysis

To reconstruct sources of the rsEEG signal, we calculated leadfield matrices based on individual brain anatomies and standard electrode positions. The T1-weighted MPRAGE images were segmented using the Freesurfer v.5.3.0 software (Fischl, 2012). We constructed a 3-shell boundary element model which was subsequently used to compute the leadfield matrix using OpenMEEG (Gramfort et al., 2010). Approximately 2,000 cortical dipolar sources were modeled for each individual. Source reconstruction was performed using exact low resolution brain electromagnetic tomography (eLORETA; Pascual-Marqui, 2007) with a regularization parameter of 0.05. We filtered the signal within the individually adjusted alpha frequency band range as well as in broadband range (3–45 Hz), squared it, and summed up across all three dipole directions. Relative alpha power (%) was then calculated in each voxel through the division of alpha power by the broadband power. The cortex surface mantle was divided into 68 regions of interest (ROIs) based on the Desikan-Killiany atlas (Desikan et al., 2006). These were further combined into five coarser ROIs (frontal, parietal, temporal, occipital, and cingulate) for the right and left hemispheres following a standard parcellation atlas (**Figure 3B)**. Relative alpha power values were averaged across each ROI.

### 2.6. Statistical Analyses

#### 2.6.1. Correlation of Age with total WMH Volume and Alpha Oscillations

Pearson correlations were calculated to examine the relationship between age and i) total or localized WMH volume (dWMH/pvWMH) and ii) the parameters of alpha oscillations in six regions at sensor space. Differences between correlations were assessed with Fisher’s r-to-z transformation implemented in R version 3.5.2 (http://www.R-project.org/). To correct for multiple comparisons, p-values were then adjusted using the False Discovery Rate (FDR; (FDR; Benjamini and Hochberg, 1995).

#### 2.6.2. Topographical Relevance Analyses of WMHs for Alpha Oscillations at Sensor Space

To identify regions in which WMHs robustly correlated with alpha oscillations, we performed whole-brain voxel-wise regressions. More precisely, we applied general linear models (GLMs) in which individual values of relative alpha power, alpha peak frequency, and LRTC were used as predictors for the topographical occurrence of WMHs, adjusting for effects of age, sex, and intracranial volume as covariates of no interest. 3D voxel-wise binary lesion maps were analyzed using *randomise* function, implemented in FSL (Winkler et al., 2014). For each statistical analysis, positive and negative contrasts were computed. The significance of results was based on threshold-free cluster enhancement (TFCE, N=10,000 permutations) with family-wise error (FWE) corrected p-values of 0.05. We further reported statistical results for the more conservative FWE threshold of p< 0.005.

#### 2.6.3. Topographical Relevance Analyses of WMHs and Alpha Power at Source Space

Since we only observed the significant results between WMHs and relative alpha power at sensor space, we implemented source-analyses only for the relative alpha power. More precisely, to assess the association between relative alpha power and whole-brain WMHs, we implemented GLMs separately for 10 ROIs with relative alpha power as a covariate of interest, and age, sex, and total intracranial volume as covariates of no interest. Because we found a positive correlation between the voxel-wise occurrence of WMHs and relative alpha power at the sensor space, we only computed a positive contrast. All statistical analyses were further corrected for multiple comparisons using TFCE based permutation testing (N=10,000) at FWE level of p< 0.05, as well as with a conservative threshold of p<0.005.

### 2.7. Sensitivity Analyses

#### 2.7.1. Control for Confounding factors

Given that different cardiovascular risk factors including body mass index (BMI), systolic blood pressure (SBP), smoking, and diabetes are associated with WMHs (Habes et al., 2016; Lampe et al., 2019; Ryu et al., 2014), we further considered these factors as potential confounders (as covariates of no interest) for the voxel-wise associations between parameters of alpha oscillations and probability of WMH occurrence in the overall sample (N=907). To assess a degree of collinearity between the regressors used in GLMs, we additionally computed variance inflation factor in R. All predictors had a variance inflation factor below 2, therefore, we concluded that models showed acceptably low multicollinearity.

#### 2.7.2. Medication

We implemented the voxel-wise inference analyses between parameters of alpha oscillations and WMHs excluding participants taking medications affecting the central nervous system (opioids, hypnotics, and sedatives, anti-parkinsonian drugs, anxiolytics, anti-psychotics, anti-epileptic drugs). The resulting sample included 801 individuals (*M=*6 8.96 ± 4.58, 323 female).

#### 2.7.3. Control Analyses

To assess the robustness of our results, we further applied voxel-wise inference analyses between the probability of WMH occurrence and absolute alpha power in the left and right occipital region at EEG source space, using age, sex, and total intracranial volume as covariates of no interest. Absolute power in both regions was log-transformed to normalize the distribution of the data for statistical analyses.

### 2.8. Mediation Analyses

We performed mediation analyses using *mediation* package (Tingley et al., 2014) in R to test the association between a predictor (X), and an outcome (Y) which can be transmitted through a mediator (M) (Hayes and Rockwood, 2017). Here, we examine whether a total or localized WMH volume (M) mediates the relationship between age as a predictor (X) and parameters of alpha oscillations at sensor space as an outcome variable (Y). Bootstrapping (n=5000) with 99% confidence intervals (CI) was used for testing the indirect effect because it does not assume normality in sampling distribution (Hayes and Rockwood, 2017). While the indirect effect shows whether age was associated with the parameters of alpha oscillations through a mediator, a total effect is the sum of indirect and direct effect. The indirect effect was considered significant if the corresponding 99% bootstrap CIs did not include zero.

### 2.9. Cognition

The Trail Making Test (TMT) is a cognitive test measuring executive function, including processing speed and mental flexibility (Reitan, 1955; Reitan and Wolfson, 1995). In the first part of the test (TMT-A) participants are asked to connect numbers in an ascending order, while in the second part (TMT-B), participants need to alternate between numbers and letters. In both TMT-A and B, the time to complete the task quantifies the performance, and lower scores indicate better performance.

We ran mediation analyses with 99% bootstrap CIs using relative alpha power in different regions as a predictor, total WMH volume as a mediator, and the task completion time in TMT-A or TMT-B as an outcome variable. The TMT data was available for 899 participants at the EEG sensor and 848 individuals at the EEG source space.

## 3. Results

### 3.1. Sample Characteristics

Details about the demographic, anthropometric, cardiovascular measures, as well as WMH volume, and alpha oscillations can be found in *Table 1*. Histograms of total WMH volume, averaged relative alpha power, its peak frequency, and LRTC across channels can be found in Supplementary Figure 2.

**Table 1.** Sample Characteristics. Abbreviations.: BMI = body mass index; DBP = diastolic blood pressure; dWMH/pvWMH = the ratio of deep/periventricular white matter hyperintensities; SD = standard deviation; SBP = systolic blood pressure; WMH = white matter hyperintensity, TMT= Trail Making Test

|  | Mean or n | Min. | Max. | SD |
| --- | --- | --- | --- | --- |
| Age (in years) | 69.49 | 60.15 | 80.03 | 4.63 |
| Female / Male | 380 / 527 |  |  |  |
| BMI (kg/m<sup>2</sup>) | 27.59 | 18.68 | 42.26 | 3.97 |
| SBP (mmHg) | 133.71 | 92.00 | 200.5 | 16.31 |
| DBP (in mmHg) | 74.54 | 43.5 | 120 | 9.06 |
| Never / former / active smokers | 517 / 319 / 71 |  |  |  |
| Diabetes (yes / no / unknown) | 143/ 748 / 16 |  |  |  |
| WMH volume (mm<sup>3</sup>) | 3935 | 127 | 78509 | 6676.76 |
| Normalized total WMH Volume | 0.0093 | 0.0003 | 0.170 | 0.015 |
| dWMH/pvWMH (%) | 0.439 | 0.011 | 3.635 | 0.402 |
| Intracranial volume (mm<sup>3</sup>) | 1729811 | 1297219 | 2466529 | 147492.5 |
| Mean relative alpha power (%) | 0.55 | 0.21 | 0.88 | 0.15 |
| Mean alpha peak frequency (Hz) | 9.4 | 7.34 | 12.01 | 0.86 |
| Mean Scaling Exponent ($\nu$) | 0.73 | 0.53 | 1.14 | 0.093 |
| TMT A (s) | 41.33 | 17.00 | 126 | 13.32 |
| TMT B (s) | 89.29 | 25.00 | 300 | 43.49 |

### 3.2. Topography and Characteristics of Alpha Oscillations

The relative alpha power at sensor space showed a maximum over the occipital channels, with a mean value of 0.66 ± 0.17 (%). Similarly, the relative alpha power at source space showed a maximum over the bilateral occipital cortex, including cuneus and lateral occipital regions with a mean value of 0.59 ± 0.18 (%). The grand-average peak frequency was 9.40 ± 0.49 Hz, showing larger values at occipital regions. The average scaling exponent (*v*) was 0.72 ± 0.017. Similarly, topographies of the scaling exponent had higher values at occipital and parietal areas as well as frontal regions (Supplementary Figure 3).

### 3.3. Association of Age with WMH Volume and Alpha Oscillations

We found a correlation between age and total WMH volume (r= 0.374, p< 0.001, Supplementary Figure 4), but not with the dWMH/pvWMH (r=0.03, p>0 .05, Supplementary Figure 5). Regarding parameters of alpha oscillations, we found that higher age was associated with decreased alpha peak frequency all EEG ROIs (r from -0.13 to -0.17, p_FDR_< 0.05), while no correlations between age and relative alpha power or LRTC were found (all p_FDR_> 0 .05). A full report of these correlations for the entire sample and by sex are provided in Supplementary Figures 6–8.

### 3.4. Topographical Association Between WMHs and Alpha Power

#### 3.4.1. Sensor Space

The voxel-wise inference analyses revealed that higher relative alpha power (%) in the frontal region was associated with higher WMH probabilities in the right body of corpus callosum ([16, -26, 32], T= 3.76, k= 653). Higher relative alpha power in the central region was associated with higher WMH probabilities in the right anterior thalamic radiation extending to the posterior corona radiata ([22, -49, 37], T= 4.44, k= 2744), while higher relative AP in the right temporal region was linked to higher WMHs in the right superior longitudinal fasciculus ([22, -49, 37], T= 4.52, k= 6893) extending to the left inferior fronto-occipital fasciculus ([-21, -53, 32], T= 4.00, k=4210). Furthermore, higher relative alpha power in the parietal region was associated with higher WMHs in the right superior corona radiata ([18, -19, 37], T= 4.05, k=4474). Similarly, for relative alpha power in the occipital region, we observed a higher prevalence of WMHs in the bilateral superior corona radiate through the body of the corpus callosum to the anterior corona radiata, including the right anterior thalamic radiation ([18, -19, 37], T= 4.39, k=9 450). Accordingly, higher voxel-wise WMH probabilities were associated with higher relative alpha power independent of age, sex, and brain size, as shown in **Figure 4** (TFCE, p<0.05, FWE-corrected). Note that using a more stringent TFCE, FWE rate of p<0.005, the correlation between the probability of WMH occurrence and relative alpha power was only evident for the occipital region ([18, -19, 37], T= 4.39, k=904). Finally, no voxel-wise associations between regional WMHs and alpha peak frequency or LRTC were observed (TFCE, p<0.05, FWE-corrected).

**Figure 4.**
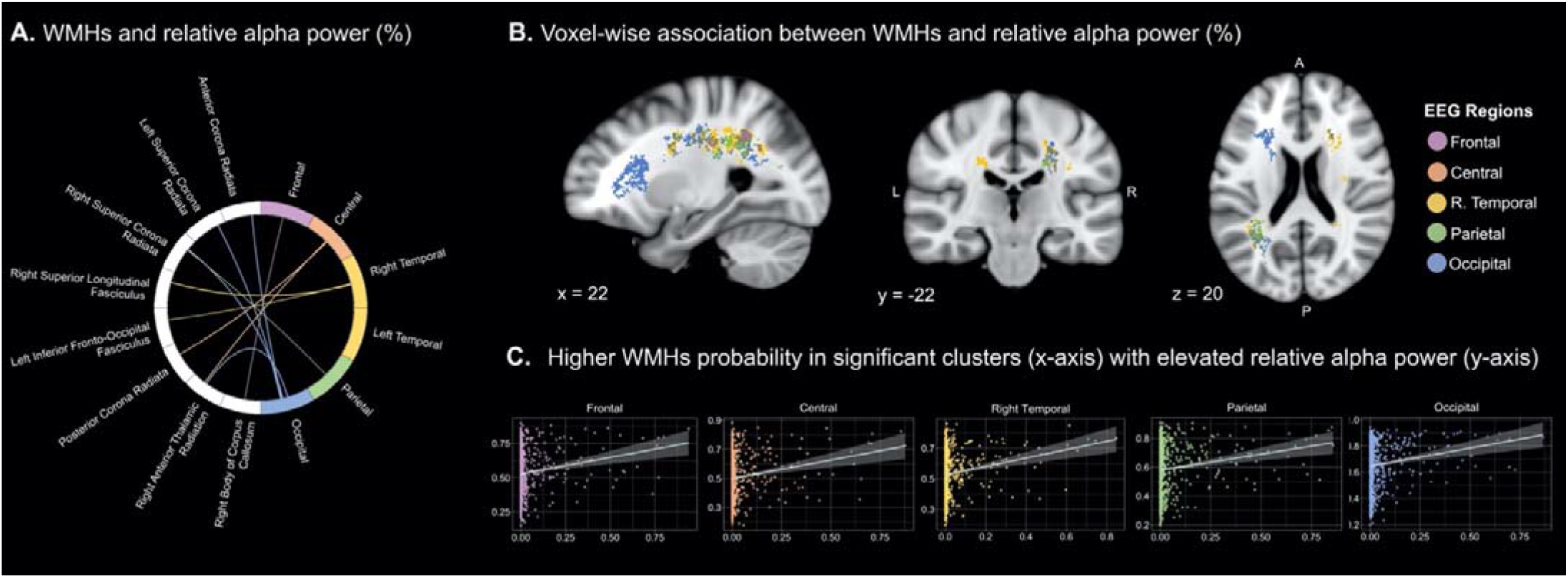
Association between white matter hyperintensities (WMHs) and relative alpha power at EEG sensor space (N=907). **A)** Schematic depiction of the significant association between regional WMHs and relative alpha power: thicker lines indicate higher t-values. **B)** We implemented nonparametric permutation testing based on whole-brain voxel-wise analysis to investigate the association between WMHs and relative alpha power (%). The brain WMH clusters show significant relation with the EEG frontal region (purple), central region (orange), right temporal region (yellow), parietal region (green), and occipital region (blue), respectively (TFCE, FWE-corrected, p<0.05 corrected for age, sex and total intracranial volume). **C)** Scatter plots show the association between WMH probability (x-axis) extracted from clusters based on significant whole-brain voxel-wise inference analyses and elevated relative alpha power (y-axis) in different EEG regions. The resulting statistical images (P-map) were further thresholded at 0.05 and binarized. Abbreviations.: A = anterior; L = left; R = right; P = posterior

#### 3.4.2. Source Space

We found that higher relative alpha power (%) in all EEG regions except for the left frontal region was associated with a higher probability of WMH occurrence (*Supplementary Table 2*, TFCE, p<0.05, FWE-corrected, Figure 5). With the stricter FWE-level of p< 0.005, the association between the occurrence of WMHs and relative alpha power was evident for left ([18, -19, 37], T= 4.29, k= 192) and right occipital regions ([18, -19, 37], T= 4.45, k=845).

**Figure 5.**
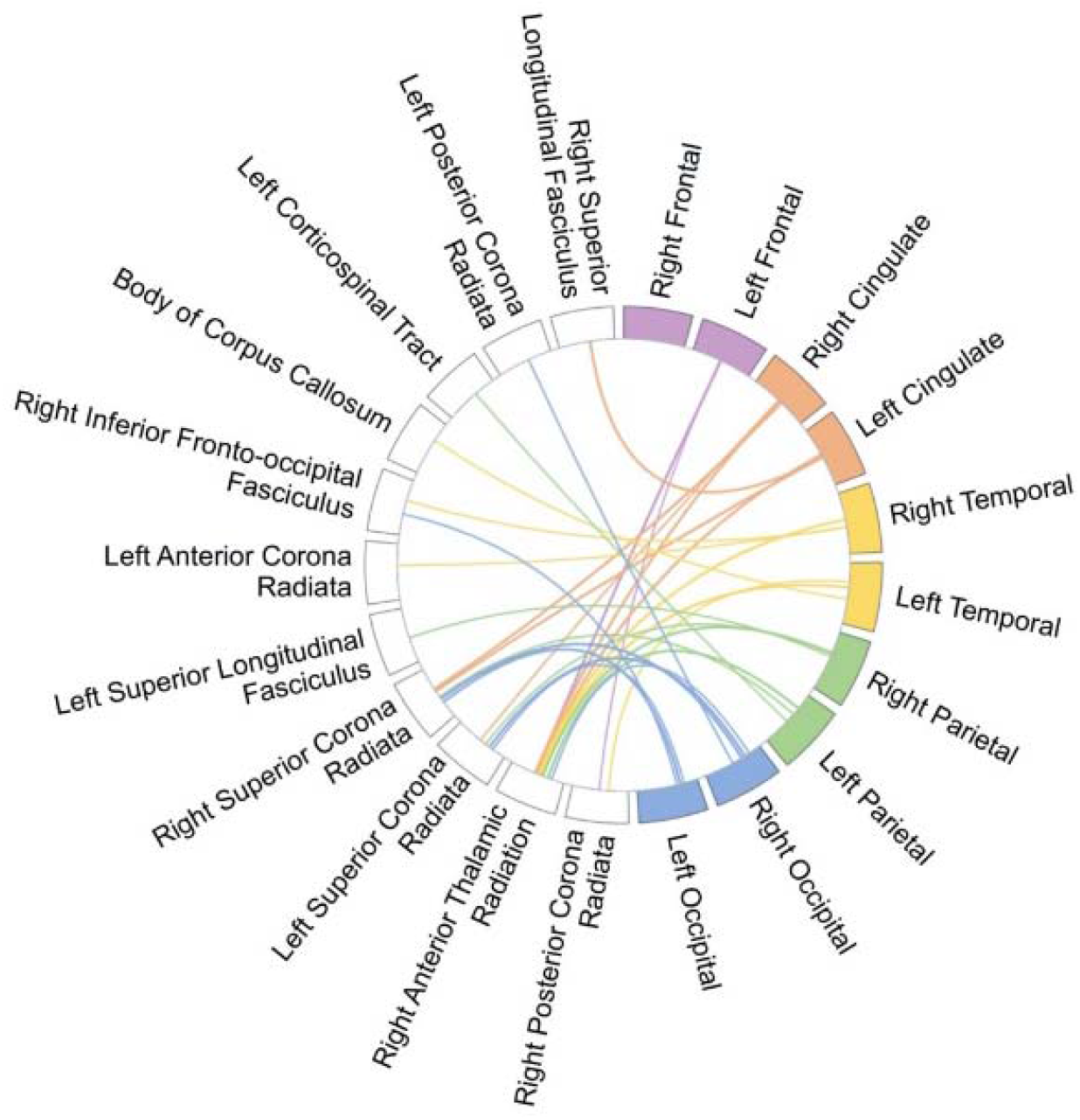
Schematic depiction of the significant association between regional WMHs and relative alpha power in EEG source space (N=855). The circular plot indicates EEG ROIs for both hemispheres at source space and their relationship to WMHs where thicker lines indicate higher t-values (See: *Supplementary Table 1*.)

### 3.5. Sensitivity Analyses

#### 3.5.1. Control for Confounding Factors

Voxel-wise inference analyses after controlling for age, sex, intracranial volume, BMI, SBP, diabetes, and smoking status yielded a similar relationship between higher WMH probability and elevated relative alpha power in the following regions: central ([22, -49, 37], T= 4.46, k= 5417), right temporal ([22, -49, 37], T= 4.52, k= 5417), left temporal ([22, -49, 37], T=4.59, k=4772), parietal ([18, -19, 37], T=3.68, k=231), and occipital ([18, -19, 37], T= 4.08, k= 4018) EEG regions across the overall sample. Note that with TFCE, FWE-corrected, p< 0.005, we did not find any clusters. Lastly, no WMH clusters were related to alpha peak frequency or LRTC (TFCE, p > 0.05, FWE-corrected).

#### 3.5.2. Medication

Voxel-wise inference analyses excluding individuals taking central nervous system medication (N=801) still indicated the association between higher prevalence of WMHs and increased relative alpha power at sensor space in the following regions: frontal ([17, 9, 31], T= 4.42, k=6880), central ([20, -30, 35], T= 4.46, k= 9063), right temporal ([20, -48, 35], T= 4.57, k=12098), left temporal ([22, -49, 37], T=4.61, k= 9408), parietal ([14, -8, 31], T= 4.61, k= 9054), and occipital ([18, -19, 37], T=4.44, k= 12,885) EEG regions. Importantly, with TFCE, FWE-corrected, p< 0.005, we identified WMHs clusters (k>2000) for occipital, left temporal, right temporal, and a small cluster (k>200) for parietal and central EEG regions. Additional voxel-wise inference analyses revealed that higher WMHs resulted in decreased alpha peak frequency in right temporal ([17, -27, 33], T= 4.00, k=138) and left temporal regions ([17, -27, 33], T=4.12, k= 503). Lastly, no WMHs clusters in the brain were related to LRTC (TFCE, p > 0.05, FWE-corrected).

#### 3.5.3. Control Analyses

Voxel-wise inference analyses with absolute alpha power similarly indicated that higher probability of WMH occurrence was associated with elevated absolute alpha power in right ([-23, 0, 36], T=3.98, k=5633) and left occipital regions ([-23, 0, 36], T= 4.05, k=5358) (TFCE, p<0.05, FWE-corrected).

### 3.6. Mediation Analyses

We examined whether total or localized (dWMH/pvWMH) WMH volume could mediate the relationship between age and relative alpha power in all cortical ROIs. Investigating the relationship between age and relative alpha power, we observed a significant indirect effect of total WMH volume in most of the cortical regions defined at sensor space (Table 2). The direct effect was not significant in any of the ROIs (99% |CI| > 0), and only in the right temporal region at sensor space did the total effect of age on relative alpha power appear to be significant (Table 2). Further, we confirmed the indirect effects of total WMH volume for relative alpha power at EEG source space for left parietal (β=0.0012, CI = [0.00006-0.002]), left (β=0.0014, CI = [0.00013-0.002]) and right occipital (β=0.0014, CI = [0.00015-0.0028]) regions. Finally, our results revealed that neither total nor localized WMH volume mediated the association of age with alpha peak frequency and LRTC at sensor space (all p>0.05).

**Table 2.** Mediation effect of total WMH volume on the association between age and relative alpha power at EEG sensor space (N=855). While the indirect or mediation effect shows whether age was associated with alpha power through a mediator (total WMH), total effect is the sum of indirect and direct effect (age on relative alpha power). The indirect effect was considered significant if the corresponding 99% bootstrap CIs did not include zero (marked in bold).

| EEG Region | frontal |  | central |  | right temporal |  | left temporal |  | parietal |  | occipital |  |
| --- | --- | --- | --- | --- | --- | --- | --- | --- | --- | --- | --- | --- |
| | $\beta$ | p or 99.5% CI | $\beta$ | p or 99.5% CI | $\beta$ | p or 99.5% CI | $\beta$ | p or 99.5% CI | $\beta$ | p or 99.5% CI | $\beta$ | p or 99.5% CI |
| Total effect c | 0.0004 | 0.742 | 0.0006 | 0.580 | **0.002** | **0.033** | 0.002 | 0.0620 | 0.0017 | 0.166 | 0.0006 | 0.584 |
| Mediation effect a*b | 0.0009 | [-0.0003, 0.0021] | 0.001 | [-0.00008, 0.0022] | **0.0013** | **[0.0003, 0.024]** | **0.0011** | **[0.00002, 0.002]** | **0.0015** | **[0.0002, 0.0028]** | **0.0014** | **[0.00012, 0.0027]** |
| Direct effect c' | -0.0005 | 0.721 | -0.0004 | 0.730 | 0.0008 | 0.44 | 0.0009 | 0.3944 | 0.0002 | 0.894 | -0.0008 | 0.557 |

### 3.7. Cognition

Compared to population-based norms (Hobert et al., 2011; Tombaugh, 2004), our sample shows similar TMT scores (Table 1), indicating good to intermediate cognitive performance. We then investigated the question of whether the relationship between relative alpha power and cognition measured by task completion time in TMT-A and B is mediated by total WMH volume. After controlling for age and sex, we found a significant indirect effect of total WMH volume on the association between TMT-A and relative alpha power only in the right temporal region (β=1.071, CI=[0.123-2.539]). In TMT-B, we observed a significant indirect effect of total WMH volume for the frontal region (β=3.399, CI = [0.252-7.896]), as shown in the Supplementary Table 2. At EEG source space, however, we did not confirm these findings. Further, in all analyses, the direct and total effects were not significant.

## 4. Discussion

The main goal of this study was to investigate whether regional WMHs affect parameters of alpha oscillations independently from age. We pursued this aim using a large sample of cognitively healthy older individuals (e.g., also based on TMT scores; Hobert et al., 2011; Tombaugh, 2004) from a population-based study (Loeffler et al., 2015). We showed distinct regional relationships between relative alpha power and WMHs: our topographical analysis suggested that higher occurrence of WMHs in superior, posterior to anterior corona radiata, as well as thalamic radiation, was related to higher relative alpha power, with strongest correlations in the bilateral occipital cortex. Adjusting for potential confounding factors including age, cardiovascular risk factors, or controlling for the effect of medication did not change these results. While the direct link between age and alpha power assessed by correlation analyses was absent, mediation analyses supported an indirect link for the existence of the relation between age and alpha power through the total WMH volume. This finding indicates why we should consider the age-related structural changes in the brain (e.g., WMHs) when we investigate the aging effects on EEG neural oscillations.

Alpha rhythm is the most salient rsEEG oscillatory phenomenon that originates from thalamocortical and cortico-cortical interactions (Bazanova and Vernon, 2014; Lopes Da Silva et al., 1997). Alterations in alpha oscillations have previously been linked to changes in different anatomical features including properties of WM (e.g., Valdés-Hernández et al., 2010). Regarding WMHs, for instance, a previous EEG-MRI study showed that higher relative alpha power in parietal regions was associated with higher scores of the prevalence of WMHs in 79 individuals with mild cognitive impairment (Babiloni et al., 2008a), consistent with our findings in this population-based sample. Previous studies with computational models have given further support for the notion that resonance properties of feedforward, cortico-thalamocortical, and intra-cortical circuits substantially influence alpha oscillations (Hindriks and van Putten, 2013). In the present study using a larger sample, we similarly observed that regional WMHs, detected mostly in superior corona radiata, containing thalamocortical fibers, affect inter-individual differences in relative alpha power. This finding was further reproduced when using alpha power values extracted from EEG source-based analysis. Although we did not observe significant association between these two measures after controlling for other confounding factors at stricter threshold (TFCE, FWE<0.005), the consistent results with regular FWE threshold at voxel-wise level suggest a possible neurophysiological link between WMHs and relative alpha power.

But, how could lesions in the WM possibly affect EEG signal which mainly reflects neural synchrony within gray matter? While in principle a hyperintensity in T2-weighted MR sequences is a quite unspecific marker of various pathologies, postmortem histopathological studies of older adults with WMHs have mostly reported demyelination, axonal loss, and other consequences of ischemic small vessel disease (Smith et al., 2000; Wardlaw et al., 2015). Myelin contributes to the speed of impulse conduction through axons, and the synchrony of impulses between distant cortical regions (Fields, 2015, 2008). Reductions of conduction velocity due to demyelination and loss of (communicating) axons are assumed to be responsible for cognitive dysfunctions which are known to be based on delicately orchestrated propagations of neuronal signals. Electrophysiologically, interactions, and synchrony between neuronal populations are reflected in rhythmic M/EEG signals, of which alpha oscillations are the most prominent ones (Bazanova and Vernon, 2014; Lopes Da Silva et al., 1997). Alpha power is a quantitative marker of the degree of synchrony in the neuronal activity of the corresponding neuronal populations (Pfurtscheller and Lopes Da Silva, 1999). While for a long-time alpha oscillations were regarded as idle rhythms of non-active brain areas, a plenitude of studies has convincingly demonstrated that alpha oscillations play an important role in many cognitive functions (Fox et al., 2016; Klimesch, 1999; Palva and Palva, 2007). For instance, in motor and sensory domains it has been shown that amplitude decreases of alpha oscillations in focal areas (i.e., reflecting cortical activation) is in turn associated with the inhibition of neighboring cortical areas (Pfurtscheller and Lopes Da Silva, 1999). This phenomenon is thought to include mutually inhibitory interactions between the chain of modules including thalamocortical and reticular nucleus neurons which are involved in the generation of alpha oscillations (Suffczynski et al., 2001). Importantly, the authors hypothesized that this surround inhibition should underlie other cognitive operations such as focused attention and stimulus selection. Such topographically specific relationships are likely to be disturbed by the alterations in conduction velocity and axonal loss in the thalamocortical circuitry (Pajevic et al., 2014). As a result of such WM disturbances, a modular organization of thalamocortical inputs and a corresponding demarcation between cortical patches of enhanced and attenuated alpha oscillations could be abolished, thus leading to a larger spread of alpha oscillations across the cortex and consequently to stronger and spatially less specific alpha oscillations. This in turn might explain a positive association between alpha power and WMHs. In addition, it is also possible to further speculate that such an elevated alpha power may result from the additional compensatory recruitment of neuronal resources to maintain an adequate brain functioning. Although we did not observe a convincing evidence for this statement in our mediation analyses involving cognition, elevated alpha power — as a consequence of WMHs — may still reflect a resilience against the cognitive decline given that cognitively healthy sample was used in the present study (e.g., Hobert et al., 2011; Tombaugh, 2004). Alternatively, the hyperactivation of alpha with WMH could also be ineffective in preserving cognitive performance or even reflect the progression of neurodegenerative alterations (Corriveau-Lecavalier et al., 2019; Pons et al., 2010).

Despite a number of reports of age-related alpha power alterations (Babiloni et al., 2006b; Lodder and van Putten, 2011; Vysata et al., 2012), in our study, we replicated other recent studies (Sahoo et al., 2020; Scally et al., 2018) which did not find strong evidence for age-related attenuations of relative alpha power. The discrepancy in findings with earlier reports could be due to the narrow age range of our participants, as well as the individually adjusted alpha frequency range based on the peak frequency. In fact, preserved peak power at peak frequency has recently been reported in an older sample (Scally et al., 2018), suggesting that any observed age-dependent power changes might be due to shifts in the frequency range at which alpha peak occurs. While our cross-sectional dataset cannot provide unequivocal evidence for a causal relationship, mediation analyses demonstrated a presence of an indirect relationship between age and alpha power through total WMHs. Currently, in the literature, there is an ongoing discussion on the interpretation and meaning of an indirect *(mediation)* effect when a total effect is not statistically significant (Hayes and Rockwood, 2017; Zhao et al., 2010). In our paper, following the suggestions by Hayes and Rockwood (2017), we also reported and interpreted the mediation effects even when a total effect was not significant. More precisely, the mediation via total WMH volume showed that higher age was associated with the elevated relative alpha power in the right temporal, parietal, and occipital regions. As mentioned before, age-related reductions of alpha power in occipital regions were previously reported in different sample populations (see detailed review: Ishii et al., 2018). As we show in this study, in healthy older adults the association between these two measures can potentially be mediated by WMH volume thus demonstrating a positive relationship between alpha power and age. Therefore, our result shows why one should potentially consider structural correlates when investigating age-related alterations in neural oscillations.

In the literature, other commonly reported age-dependent changes in spectral parameters of EEG include slowing of the alpha peak (Knyazeva et al., 2018). We replicated the slowing of the alpha peak frequency with increasing age despite the narrow age range. Alpha peak slowing has previously been suggested to be linked to a less efficient coordination of neuronal activity in this frequency range (Mierau et al., 2017). We further explored the relationship between age and LRTC in the amplitude envelope of alpha oscillations that capture scale-free dynamics of resting-state oscillations. LRTC has previously been linked to the presence of a critical state in neural networks, which is characterized by the balance of excitation and inhibition (Poil et al., 2012) that has been suggested to be optimal for the processing of information in the human brain. Regarding the association between age and LRTC, previous studies have shown that the observed age-related changes might be dependent on age range — it increases from childhood to early adulthood, after which it stabilizes (Nikulin and Brismar, 2005; Smit et al., 2011). In accordance with these previous findings, in our sample of older adults, we observed no pronounced age-related LRTC attenuations. The fact that WMHs were correlated with alpha power but not with LRTC indicates that temporal dynamics have more flexibility in adjusting to white matter lesions since they are largely based on cortico-cortical interactions which are not reflected in WMH (Beggs and Plenz, 2003).

## 5. Limitations

While a strength of this study is the large population-based sample, the study design is cross-sectional and does not allow making inferences about the directionality of the association between WMHs and alpha oscillations. Longitudinal studies are required to further clarify these associations. Research using other advanced techniques such as quantitative MRI or specific assessment of tissue properties with ultra-high field MRI combined with intracranial EEG recording could further provide valuable insights into the nature of the relationship between WM properties and alpha oscillations. We performed a relatively coarse parcellation of the brain at EEG source space analysis due to the relatively small number of electrodes (n= 31). A denser spatial sampling of the EEG (not available in the present cohort) would allow investigation of this relationship with better spatial precision. Finally, while our study aimed to investigate the effect of WMHs on properties of alpha oscillations, future research on aging using microstructural integrity assessed by DTI would benefit from additional connectivity-based measures including phase synchrony (Hinault et al., 2020; Quandt et al., 2020).

## 6. Conclusion

Using sensitive high-resolution neuroimaging techniques in cognitively healthy older adults (N=907), we showed that elevated relative alpha power is related to a higher probability of WMHs, supporting the idea that an elevated alpha power as a consequence of WMHs may result from the additional recruitment of compensatory neuronal resources during aging. Importantly, our study provides evidence that the changes in alpha oscillations do not relate to aging *per se* but rather depend on the impact of age-related neuropathology, such as WMHs. Our findings thus suggest that longitudinal EEG recordings might be sensitive for the detection of alterations in neuronal activities due to progressive structural changes in WM.

## Acknowledgments

We thank all members of the Leipzig Research Center for Civilization Diseases (LIFE) study center for conducting the LIFE-Adult study and also all participants for their valuable collaboration.

## 8. Data availability

Anonymized data will be made available upon request through the application procedure carried out by the LIFE-Study administration (https://life.uni-leipzig.de/de/erwachsenen_kohorten/life_adult.html).

## Supplementary Material

**Supplementary Figure 1.**
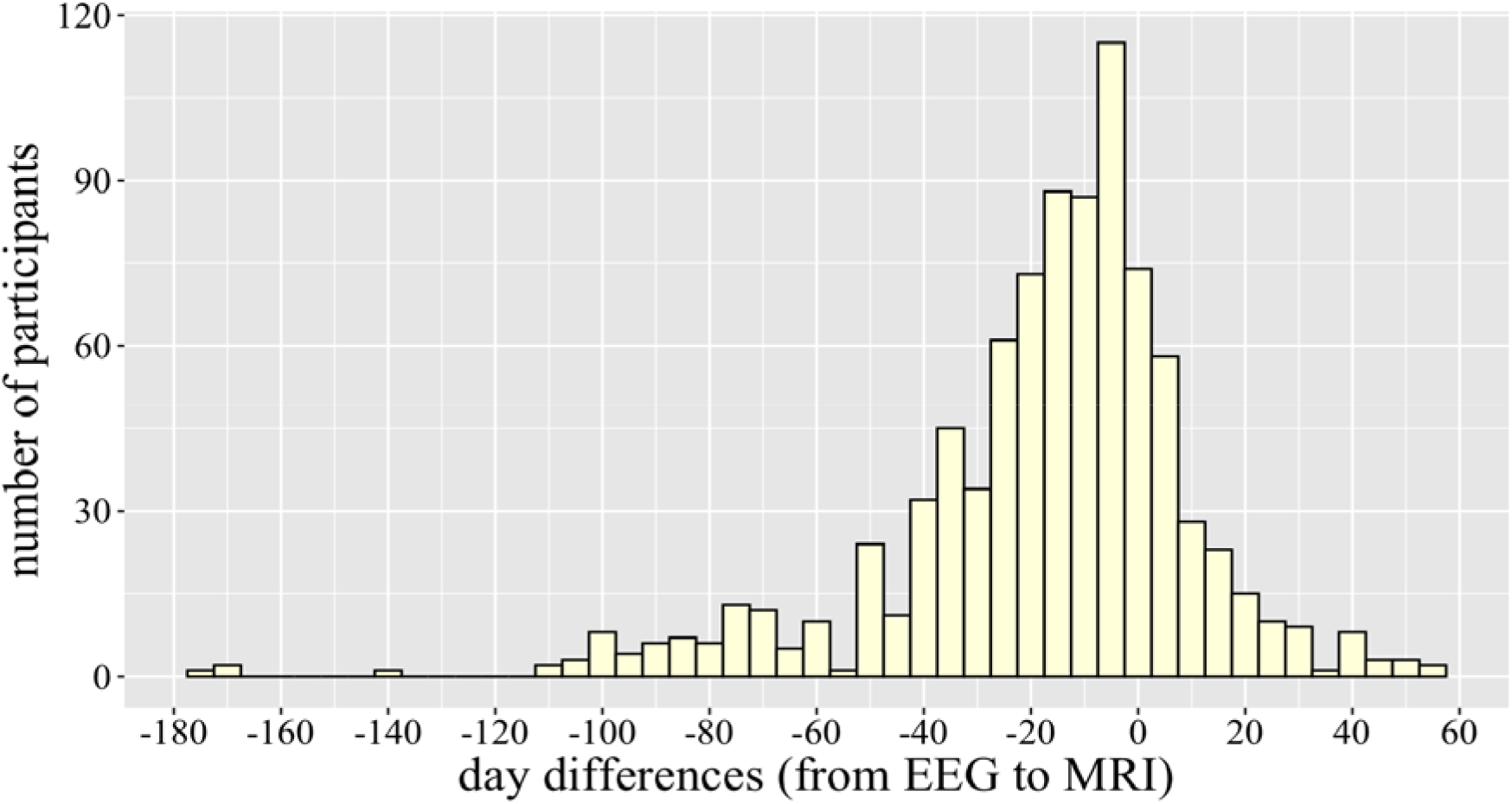
Histogram of the day differences between EEG and MRI acquisition points. While the averaged (absolute) day difference across participants was 23.4 days, the minimum day was 0, maximum was 175 days. All variables in the supplementary Figure 2 are presented as mean (M) ± standard deviation (SD). Before the statistical analyses, we used the Box-Cox method (λ value) (Sakia, 1992) to determine the type transformation on the parameters of alpha oscillations. Since the majority of the variables after the necessary transformation did not pass Shapiro-Wilk normality tests at the 0.05 significance level, we decided to keep the original values.

**Supplementary Figure 2.**
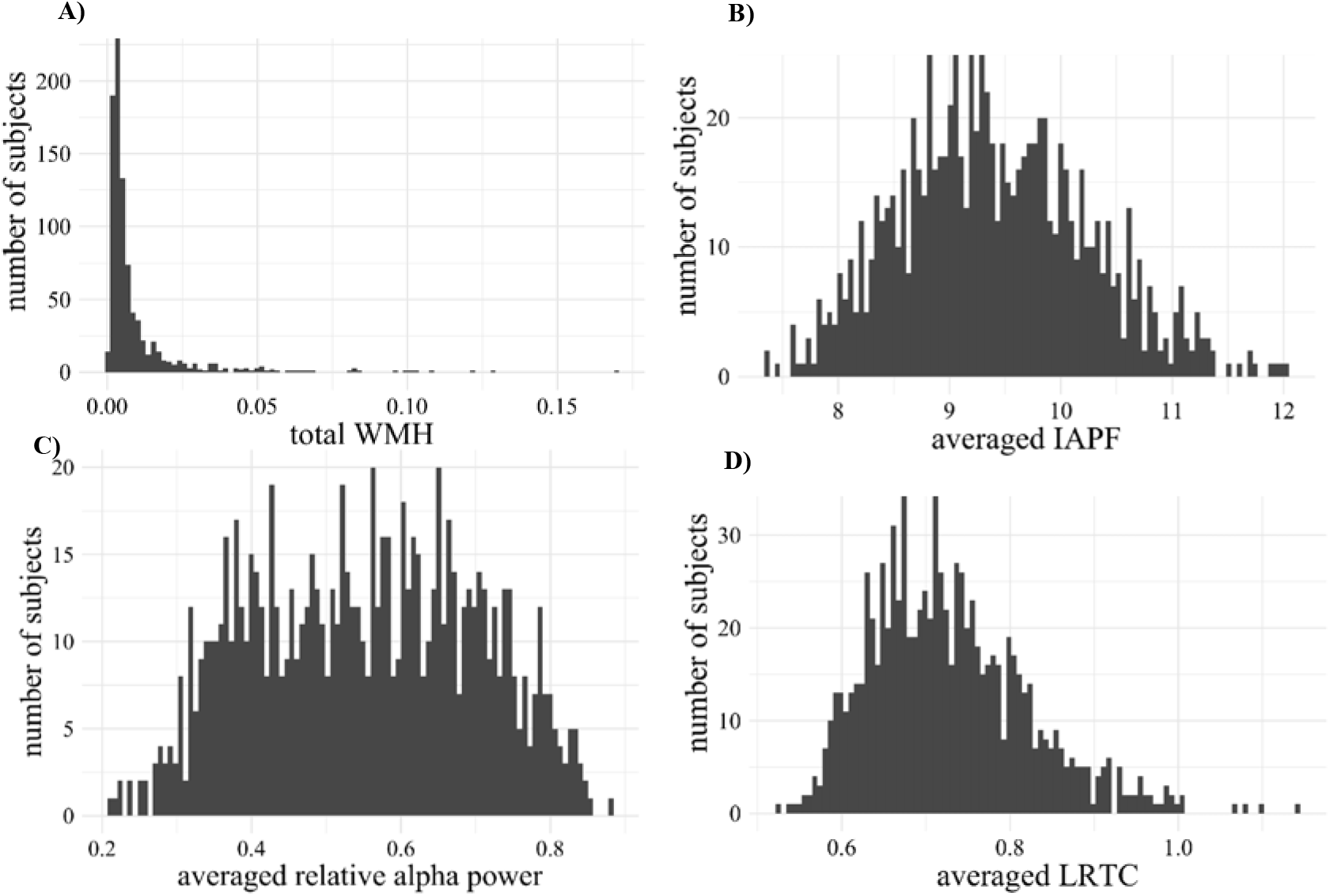
The four histograms show the distribution of **A)** normalized total white matter hyperintensity (WMH), **B)** individual alpha peak frequency (IAPF), **C)** relative alpha power, and **D)** long-range temporal correlation (LRTC) averaged across 31 EEG channels. Note that total WMH volume further normalized to head size by total intracranial volume and log-transformed.

**Supplementary Figure 3.**
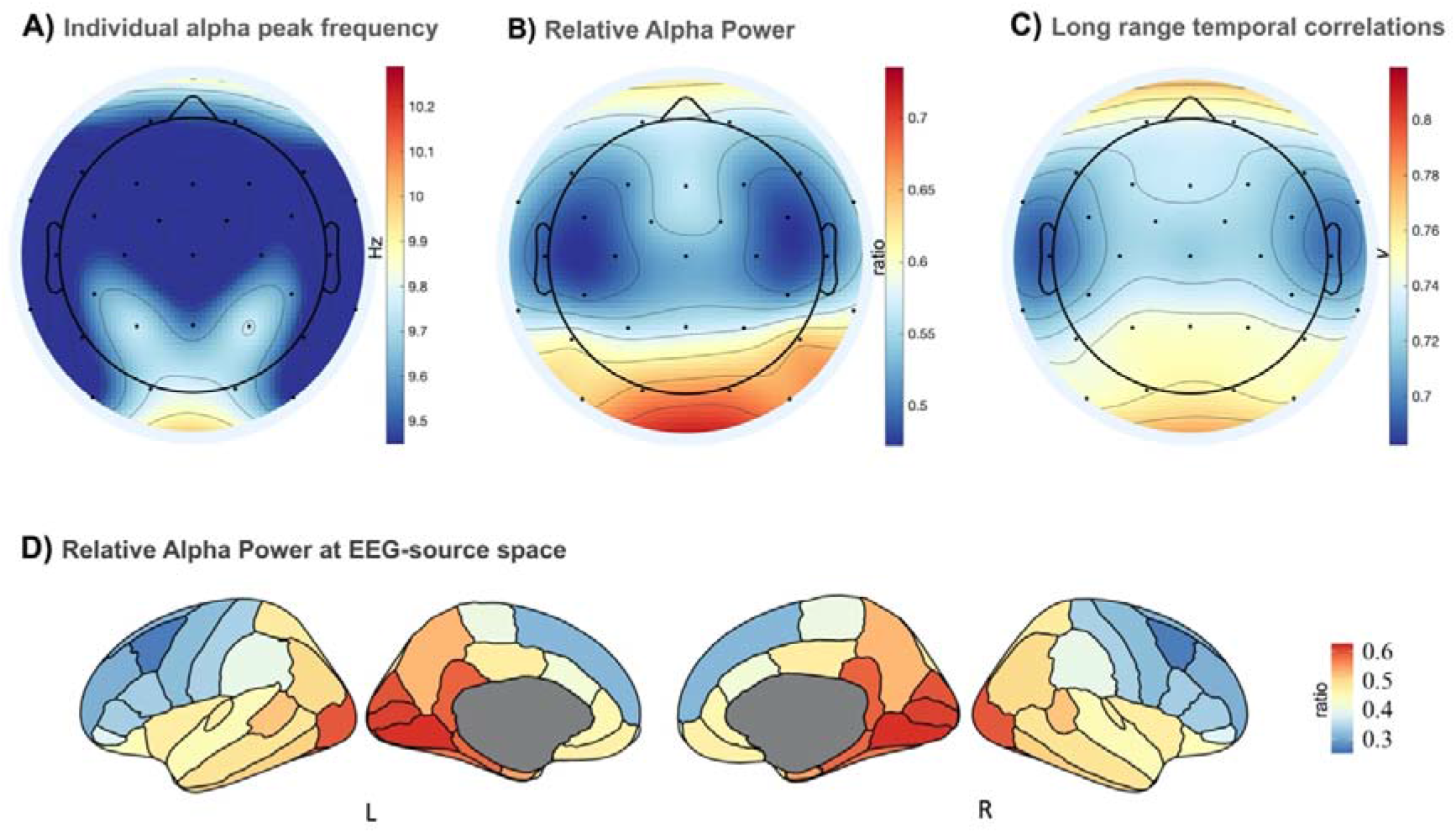
Grand-average topographic maps of alpha band measures in EEG. **A)** Individual alpha peak frequency (Hz); **B)** Relative alpha power (%); **C)** Long-range temporal correlations (*v*). **D)** Grand-average of relative alpha power at EEG source space across 68 regions based on Desikan-Killiany Atlas.

**Supplementary Figure 4.**
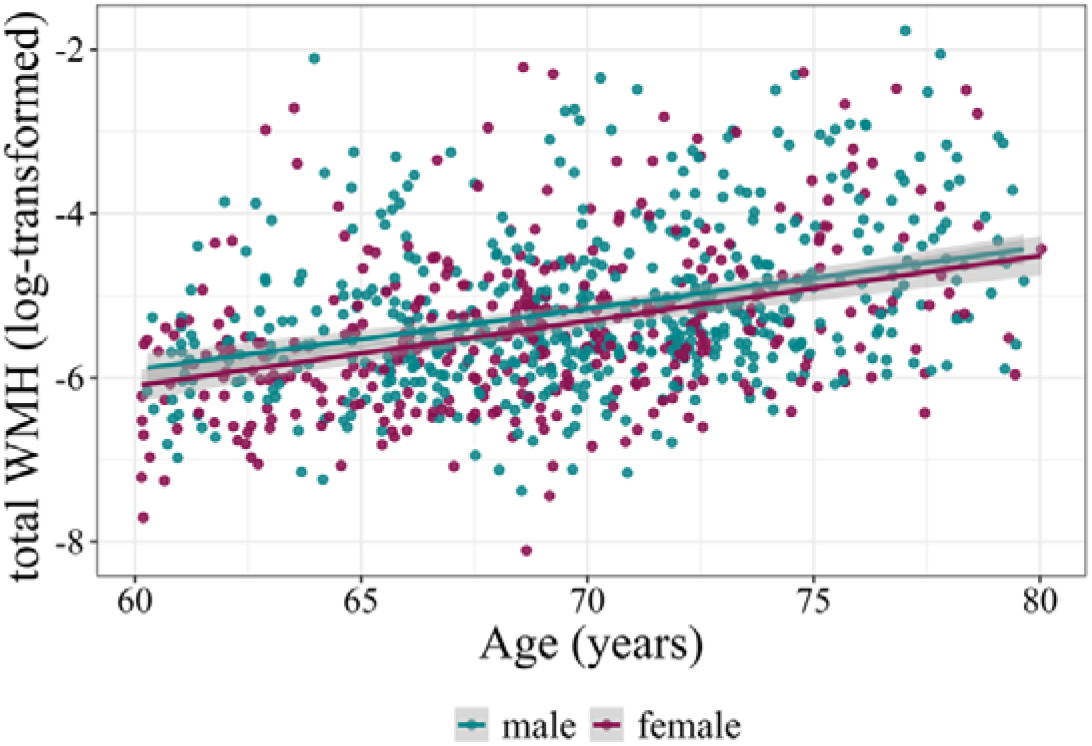
Association between age (x-axis) and total white matter hyperintensity (WMH, y-axis) in LIFE-Adult sample (N=907). There was a significant correlation between age and total WMH (r=0.374, p<0.001 in all; r=0.376, p<0.001 in females; r=0.355, p<0.001 in males). Note that Total WMH volume further normalized by total intracranial volume and log-transformed.

**Supplementary Figure 5.**
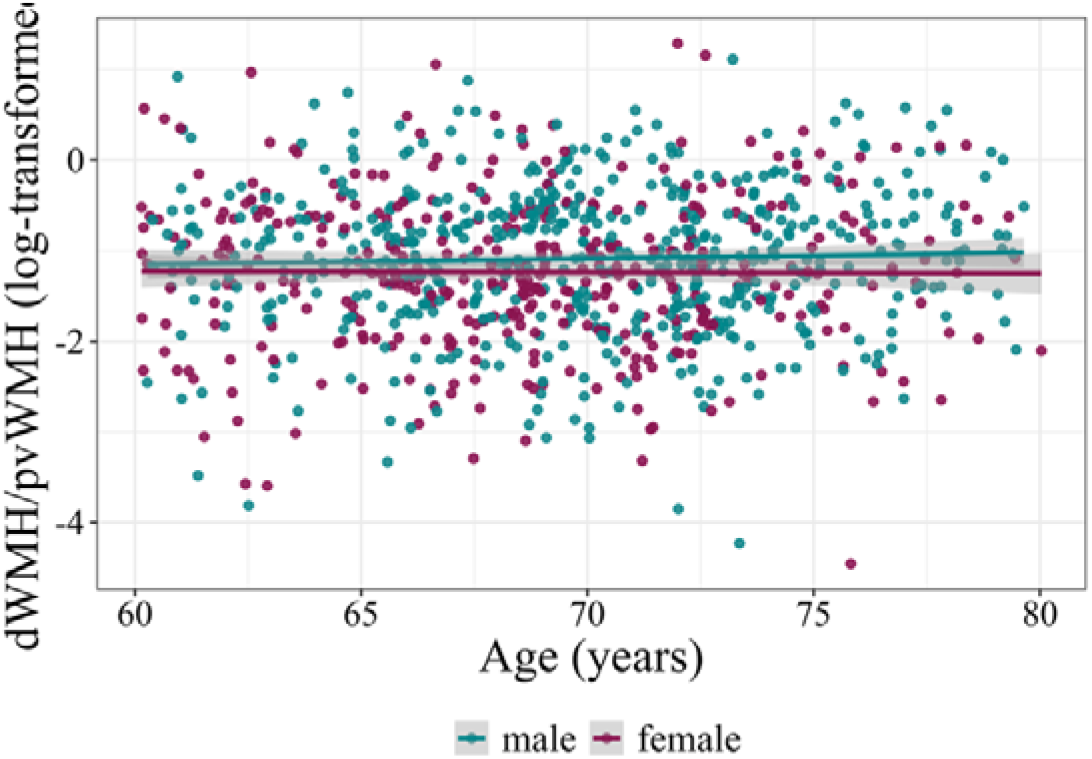
Association between age (x-axis) and regional white matter hyperintensity as the ratio of deep WMH and periventricular WMH (y-axis) in LIFE-Adult sample (N=907) (r=0.03, p=0.354 in all; r=-0.005, p=0.912 in females; r=0.038, p =0.379 in males)

**Supplementary Figure 6.**
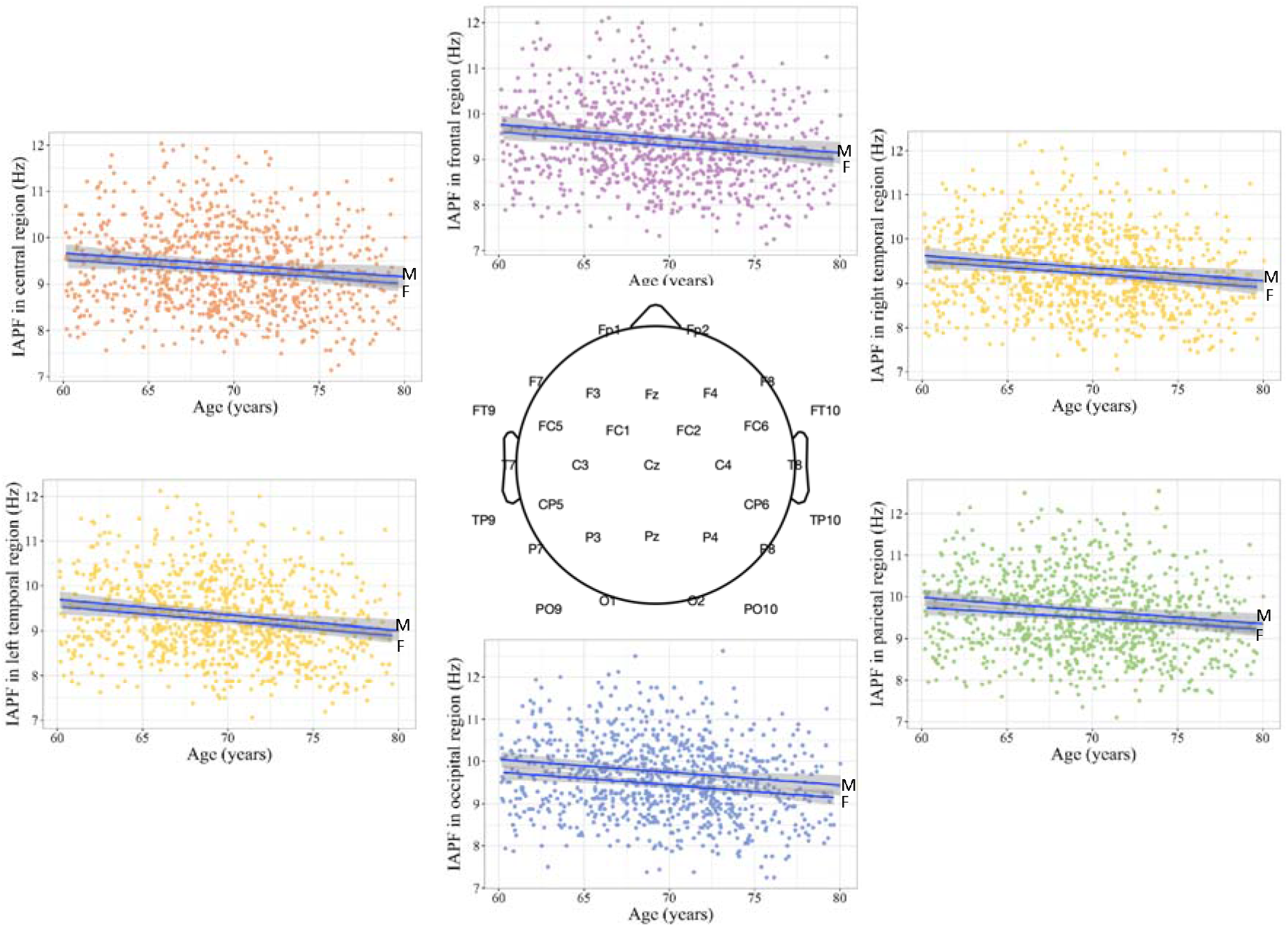
Association between age (x-axis, in years) and individual alpha peak frequency (IAPF, y-axis, in Hz) in EEG different regions. The correlations between two measures were not significant after FDR correction and none of the pairwise correlations differed from each other. Abbr.: F-female, M-male - Frontal (r=-0.16, p<0.001 in all, r=-0.147, p=0.004 in females, r=-0.16, p=0.0001 in males)
- Central (r=-0.130, p<0.001 in all, r=-0.12, p=0.01 in females, r=-0.12, p=0.004 in males)
- Left temporal (r=-0.17, p<0.001 in all, r=-0.166, p=0.001 in females, r=-0.168, p=0.0001 in males)
- Right Temporal (r=-0.156, p<0.001 in all, r=-0.141 p=0.006 in females; r=-0.146, p=0.0009 in males)
- Parietal (r=-0.158, p<0.001 in all, r=-0.144 p=0.005 in females; r=-0.143, p=0.001 in males)
- Occipital (r=-0.170, p<0.001 in all, r=-0.13, p=0.01 in females, r=-0.164, p=0.0001 in males)

**Supplementary Figure 7.**
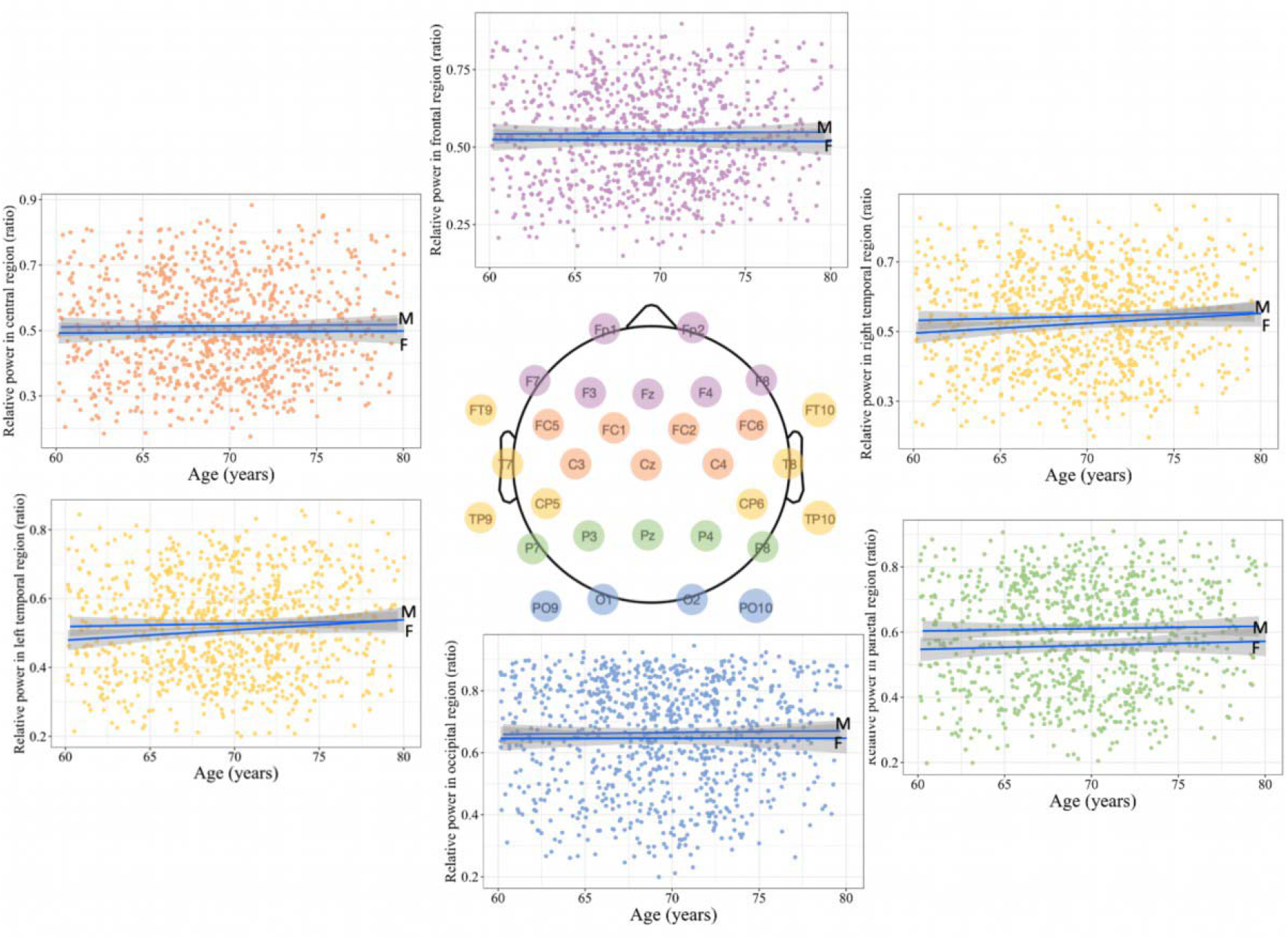
Association between age (x-axis, in years) and relative alpha power (y-axis, ratio expressed in %) in different EEG regions. The correlations between two measures were not significant after FDR correction and none of the pairwise correlations differed from each other. Abbr.: F-female, M-male - Frontal (r=0.010, p=0.742 in all, r=0.008, p=0.868 in females, r=-0.008, p=0.837 in males)
- Central (r=0.009, p=0.850 in all, r=0.012, p=0.781 in females, r=0.019, p=0.565 in males)
- Left temporal (r=0.0065, p=0.048 in all; r=0.098, p=0.056 in females, r=0.027, p=0.52 in males)
- Right Temporal (r=0.071, p=0.03 in all, r=0.090, p=0.07 in females; r=0.040, p=0.355 in males)
- Parietal (r=0.04, p=0.16 in all, r=0.033, p=0.51 in females, r=0.02, p =0.62 in males)
- Occipital (r=0.016, p=0.61 in all, r=0.001, p=0.98 in females, r=0.016, p=0.69 in males)

**Supplementary Figure 8.**
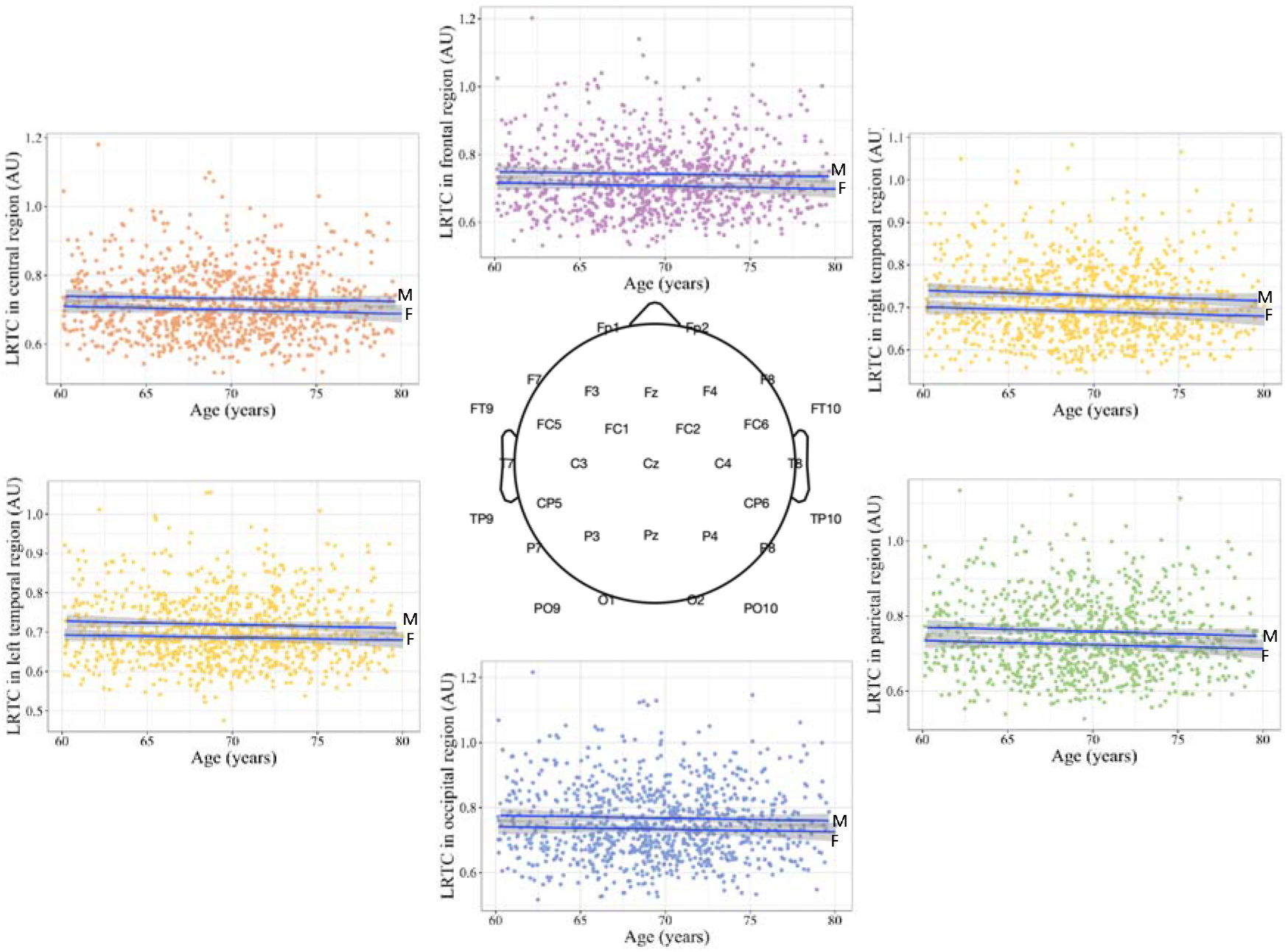
Association between age (x-axis, in years) and scaling exponent (*v*) for long-range temporal correlations LRTC, y-axis) in different EEG regions (represented in different colors). The correlations between two measures were not significant after FDR correction. - Frontal (r=0.02, p=0.540 in all, r=0.04, p=0.409 in females, r=-0.04, p=0.312 in males)
- Central (r=0.04, p=0.288 in all, r=0.07, p=0.166 in females, r=0.05, p=0.192 in males)
- Left temporal (r=0.05, p=0.109 in all; r=0.098, p=0.07 in females, r=0.07, p=0.09 in males)
- Right Temporal (r=0.071, p=0.03 in all, r=0.08, p=0.112 in females; r=0.040, p=0.355 in males)
- Parietal (r=0.04, p=0.248 in all, r=0.06, p=0.284 in females, r=0.06, p =0.127 in males)
- Occipital (r=0.023, p=0.513 in all, r=0.03, p=0.558 in females, r=0.05, p=0.252 in males)

**Supplementary Table 1.** Positive correlation between the probability of white matter hyperintensity (WMH) occurrence and relative alpha power (%) at EEG source space. Peak voxel MNI coordinates (x, y, z) and cluster size (k) for the association between WMHs probability and relative alpha power for five regions of interest for each hemisphere at source space across 855 older adults (TFCE, p < 0.05, FWE-corrected).

| EEG Region | MRI Region | x | y | z | k | T-value |
| --- | --- | --- | --- | --- | --- | --- |
| Left Frontal | Right Posterior Corona Radiata /<br>Right Anterior Thalamic Radiation | 21 | -46 | 36 | 219 | 4.38 |
| Right Cingulate | Right Anterior Thalamic Radiation /<br>Right Anterior Thalamic Radiation | 22 | -49 | 37 | 2310 | 4.33 |
|  | Left Superior Corona Radiata | -22 | 6 | 31 | 655 | 4.29 |
|  | Right Superior Corona Radiata | 29 | -46 | 26 | 359 | 3.65 |
| Left Cingulate | Right Anterior Thalamic Radiation /<br>Right superior Longitudinal Fasciculus | 22 | -49 | 37 | 3280 | 4.44 |
|  | Left Superior Corona Radiata | -22 | 6 | 31 | 597 | 4.33 |
| Right Temporal | Right Anterior Thalamic Radiation | 20 | -50 | 36 | 4669 | 4.57 |
|  | Left Anterior Corona Radiata | -18 | 18 | 27 | 2044 | 4.14 |
|  | Right Inferior Fronto-occipital Fasciculus | 34 | -49 | 0 | 129 | 3.68 |
| Left Temporal | Right Anterior Thalamic Radiation | 20 | -50 | 36 | 602 | 4.63 |
|  | Body of Corpus Callosum | 16 | -5 | 36 | 279 | 3.63 |
|  | Right Posterior Corona Radiata | 19 | -30 | 35 | 132 | 4.13 |
| Right Parietal | Right Anterior Thalamic Radiation | 20 | -50 | 36 | 3983 | 4.72 |
|  | Left Superior Corona Radiata | -19 | 11 | 28 | 824 | 3.98 |
|  | Left Superior Longitudinal Fasciculus | -24 | -12 | 40 | 210 | 4.12 |
| Left Parietal | Right Superior Corona Radiata/Left<br>Corticospinal Tract | 19 | -25 | 36 | 634 | 3.91 |
|  | Right Anterior Thalamic Radiation | 20 | -50 | 36 | 618 | 4.75 |
| Right Occipital | Right Superior Corona Radiata | 18 | -19 | 37 | 8339 | 4.45 |
|  | Left Superior Corona Radiata | -19 | 9 | 29 | 1070 | 4.41 |
|  | Left Posterior Corona Radiata/Anterior<br>Thalamic Radiation | -24 | -27 | 31 | 100 | 3.94 |
| Left Occipital | Right Superior Corona Radiata | 18 | -19 | 37 | 7304 | 4.29 |
|  | Left Superior Corona Radiata | -19 | 9 | 29 | 450 | 4.19 |
|  | Right Inferior Fronto-occipital Fasciculus | 34 | -37 | -4 | 175 | 3.94 |
|  | Left Superior Corona Radiata | -20 | -6 | 32 | 133 | 3.66 |

**Supplementary Table 2.** Mediation effect of total WMH volume on the association between relative alpha power at EEG sensor space and cognition, measured by trail making test (TMT), corrected by age and sex. While the indirect or mediation effect shows whether relative alpha power was associated with TMT-A or -B through a mediator (total WMH volume), total effect is the sum of indirect and direct effect (relative alpha power on TMT-A or TMT-B). The indirect effect was considered significant if the corresponding 99% bootstrap CIs did not include zero, marked in bold.

| Cognition | EEG Region | frontal |  | central |  | right temporal |  | left temporal |  | parietal |  | occipital |  |
| --- | --- | --- | --- | --- | --- | --- | --- | --- | --- | --- | --- | --- | --- |
| | | $\beta$ | p or 99.5% CI | $\beta$ | p or 99.5% CI | $\beta$ | p or 99.5% CI | $\beta$ | p or 99.5% CI | $\beta$ | p or 99.5% CI | $\beta$ | p or 99.5% CI |
| TMT-A | Total effect c | -0.739 | 0.81 | -1.658 | 0.56 | 0.891 | 0.7828 | -0.669 | 0.85 | -1.180 | 0.65 | -3.596 | 0.15 |
|  | Mediation effect a*b | 0.270 | [-0.333, 1.133] | 0.236 | [-0.362, 1.07] | **1.071** | **[0.123, 2.539]** | 0.270 | [-0.357, 1.24] | 0.160 | [-0.25, 0.76] | 0.280 | [-0.307, 1.111] |
|  | Direct effect c' | -1.009 | 0.74 | -1.894 | 0.51 | -0.181 | 0.956 | -0.939 | 0.78 | -1.020 | 0.70 | -3.876 | 0.12 |
| TMT-B | Total effect c | 12.741 | 0.157 | 8.763 | 0.349 | 7.519 | 0.435 | 8.808 | 0.380 | 7.446 | 0.40 | 7.897 | 0.308 |
|  | Mediation effect a*b | **3.399** | **[0.252, 7.896]** | 2.978 | [-0.703, 7.472] | 2.879 | [-0.055, 7.123] | 2.206 | [-0.285, 6.218] | 2.064 | [-0.14, 6.38] | 2.072 | [-0.02, 5.707] |
|  | Direct effect c' | 9.342 | 0.300 | 5.785 | 0.540 | 4.639 | 0.626 | 6.602 | 0.518 | 5.382 | 0.53 | 5.824 | 0.456 |

